# Changes in mitochondrial distribution occur at the axon initial segment in association with neurodegeneration in *Drosophila*

**DOI:** 10.1101/2024.02.14.580288

**Authors:** Andrew P. K. Wodrich, Brent T. Harris, Edward Giniger

## Abstract

Changes in mitochondrial distribution are a feature of numerous age-related neurodegenerative diseases. In *Drosophila*, reducing the activity of Cdk5 causes a neurodegenerative phenotype and is known to affect several mitochondrial properties. Therefore, we investigated whether alterations of mitochondrial distribution are involved in Cdk5-associated neurodegeneration. We find that reducing Cdk5 activity does not alter the balance of mitochondrial localization to the somatodendritic vs. axonal neuronal compartments of the mushroom body, the learning and memory center of the *Drosophila* brain. We do, however, observe changes in mitochondrial distribution at the axon initial segment (AIS), a neuronal compartment located in the proximal axon involved in neuronal polarization and action potential initiation. Specifically, we observe that mitochondria are partially excluded from the AIS in wild-type neurons, but that this exclusion is lost upon reduction of Cdk5 activity, concomitant with the shrinkage of the AIS domain that is known to occur in this condition. This mitochondrial redistribution into the AIS is not likely due to the shortening of the AIS domain itself but rather due to altered Cdk5 activity. Furthermore, mitochondrial redistribution into the AIS is unlikely to be an early driver of neurodegeneration in the context of reduced Cdk5 activity.

**Summary statement:** In *Drosophila*, mitochondria are excluded from the axon iniCal segment, a neuronal compartment that regulates neuron polarity and axon potenCals, and this paYern is disrupted in a model of neurodegeneraCon.

## 1. INTRODUCTION

Mitochondrial changes have been linked to both aging and age-related neurodegenerative disease (Lezi & Swerdlow, 2012; López-Otin et al., 2023). One property of mitochondria that influences cellular health is their subcellular distribution, which is especially important in highly polarized cells such as neurons (Misgeld & Schwarz, 2017). Indeed, abnormal mitochondrial distribution within neurons is associated with aging and multiple neurological disorders (Bereiter-Hahn, 2014; Iijima-Ando et al., 2009; Pigino et al., 2003; Stommel et al., 2007; Vagnoni et al., 2016).

Mitochondrial distribution within neurons is controlled, in part, by the axon initial segment (AIS). The AIS is a neuronal compartment located in the proximal axon between the axonal and somatodendritic compartments (Huang & Rasband, 2018). The AIS is organized in both vertebrates and invertebrates by large ankyrin proteins that serve as scaffolding to link voltage-gated sodium and potassium channels to the underlying cytoskeletal network (Eichel et al., 2022; Jegla et al., 2016; Leterrier, 2018; Trunova et al., 2011). The dense cluster of voltage-gated ion channels in the AIS facilitates action potential initiation and makes the AIS a critical domain governing the function of the entire neuron (Huang & Rasband, 2018). Additionally, the structural proteins of the AIS, including actin, ankyrins, β-spectrins, and microtubule-associated proteins, maintain neuronal polarity by regulating the passage of both molecules and organelles, including mitochondria, through the AIS (Huang & Rasband, 2018; Jones & Svitkina, 2016; Nirschl et al., 2017). Certain axonal and somatodendritic proteins, moreover, are selectively excluded from accumulating in the AIS itself (Huang & Rasband, 2018; Jones & Svitkina, 2016). Importantly, the size, structure, and function of the AIS are all known to change in the context of aging and nervous system pathology (Atapour & Rosa, 2017; Cruz et al., 2009; Jahan et al., 2022; Marin et al., 2016).

Cdk5 is a non-canonical cyclin-dependent kinase associated with age-related neurodegenerative diseases (Pao & Tsai, 2021). The activity of Cdk5 is restricted to post-mitotic neurons due to the regulated expression of its activating subunit (Pao & Tsai, 2021). In mammals, Cdk5 is activated by binding to the proteins p35 or p39, while in *Drosophila* Cdk5 has only a single activator, Cdk5α (also called D-p35), that is orthologous to the mammalian activating subunits (Connell-Crowley et al., 2000; Tang et al., 1995; Tsai et al., 1994). Among its many functions, Cdk5 is a major tau kinase, and alterations of its activity are associated with age-related neurodegenerative diseases such as Alzheimer disease, Parkinson disease, and amyotrophic lateral sclerosis in humans and mammalian model systems (Alvira et al., 2008; Bajaj et al., 1998; Nguyen et al., 2001; Patrick et al., 1999; Qu et al., 2007; Smith et al., 2003). Reducing the activity of Cdk5 in *Drosophila* by knocking out Cdk5α (hereafter: Cdk5α-KO or KO) is sufficient to mimic many of the cellular and organismal features of mammalian neurodegenerative disease. Specifically, Cdk5α-KO flies display altered innate immunity, impaired autophagy, locomotor impairment, and shortened lifespan (Connell-Crowley et al., 2007; Howell et al., 2012; Kissler et al., 2009; Nandi et al., 2017; Shukla et al., 2019; Spurrier et al., 2018). These flies also show adult-onset, age-dependent degeneration of the mushroom body (MB), the learning and memory center of the fly brain, as well as degeneration of dopaminergic neurons (Shukla et al., 2019; Spurrier et al., 2018; Trunova & Giniger, 2012).

Altering Cdk5 activity also affects the AIS. Specifically, it has been demonstrated that Cdk5α-KO in *Drosophila* causes AIS shrinkage or absence in MB neurons, followed by local swelling of the AIS and subsequent axonal degeneration (Spurrier et al., 2019; Trunova et al., 2011). Moreover, expressing a dominant-interfering fragment of the *Drosophila* neuronal ankyrin, Ank2-L4, induces similar structural disruption and shrinkage of the AIS, and this is also sufficient to cause axonal degeneration, neuron cell loss, and increased mortality (Higham et al., 2019; Jegla et al., 2016; Spurrier et al., 2019). Genetic experiments, however, suggested that the mechanism of AIS disruption by Ank2-L4 expression is distinct from that induced by Cdk5α-KO (Spurrier et al., 2019).

Previous transcriptional profiling has suggested that altered mitochondrial properties play a role in the pathology of many neurodegenerative processes, including Cdk5-associated, age-related neurodegeneration (Spurrier et al., 2018). Therefore, we sought to investigate whether Cdk5-associated perturbations of mitochondrial properties, including mitochondrial distribution, might be involved in Cdk5-associated neurodegeneration.

Here, we show that reducing Cdk5 activity by Cdk5α-KO does not disrupt overall mitochondrial distribution between the axonal and somatodendritic compartments, suggesting that mitochondrial localization deficits to these compartments are not a part of the mechanism of Cdk5-associated neurodegeneration. However, Cdk5α-KO does cause a redistribution of mitochondria into the AIS, a domain that we show limits or excludes accumulation of mitochondria in both healthy larval and adult MB neurons. Our data suggest that this redistribution of mitochondria into the AIS likely occurs because of altered Cdk5 activity rather than simply as a consequence of disruption of the AIS *per se*. Finally, while our findings indicate the redistribution of mitochondria into the AIS correlates with axonal degeneration in two different genetic scenarios, our data suggest that the mitochondrial redistribution into the AIS alone is unlikely to serve as the primary trigger for initiating axonal degeneration.

## 2. RESULTS

### 2.1. Cdk5α-KO does not cause gross disruption of mitochondrial distribution between somatodendritic and axonal compartments of larval and adult MB neurons

To determine if reducing Cdk5 activity grossly disrupts mitochondrial distribution within neurons across the lifespan, we expressed a mitochondrially-tagged fluorophore (*UAS-dsRed.mito 47A*) in the *Drosophila* MB using *201Y-GAL4*. The MB was selected for analysis because this region of the *Drosophila* central brain has a stereotyped neuronal architecture with clearly separable axonal and somatodendritic subcellular compartments (Fig. 1A), and because neurodegeneration is readily apparent in this structure upon loss of Cdk5 function by Cdk5α-KO, as assayed both by axonal fragmentation and neuronal cell death (Spurrier et al., 2018). To compare the relative distribution of mitochondria within neurons, a ratio was calculated of the mitochondrial fluorophore intensity in the somatodendritic compartment to that of the axonal compartment of MB neurons (Fig. 1B). Relative to wild-type controls (hereafter: WT), Cdk5α-KO does not alter the ratio of somatodendritic to axonal mitochondria in 3^rd^ instar larvae, 10 day-old adults, nor even in 30 day-old adults, a stage at which degeneration is apparent (Fig. 1C-I).

**Figure 1.**
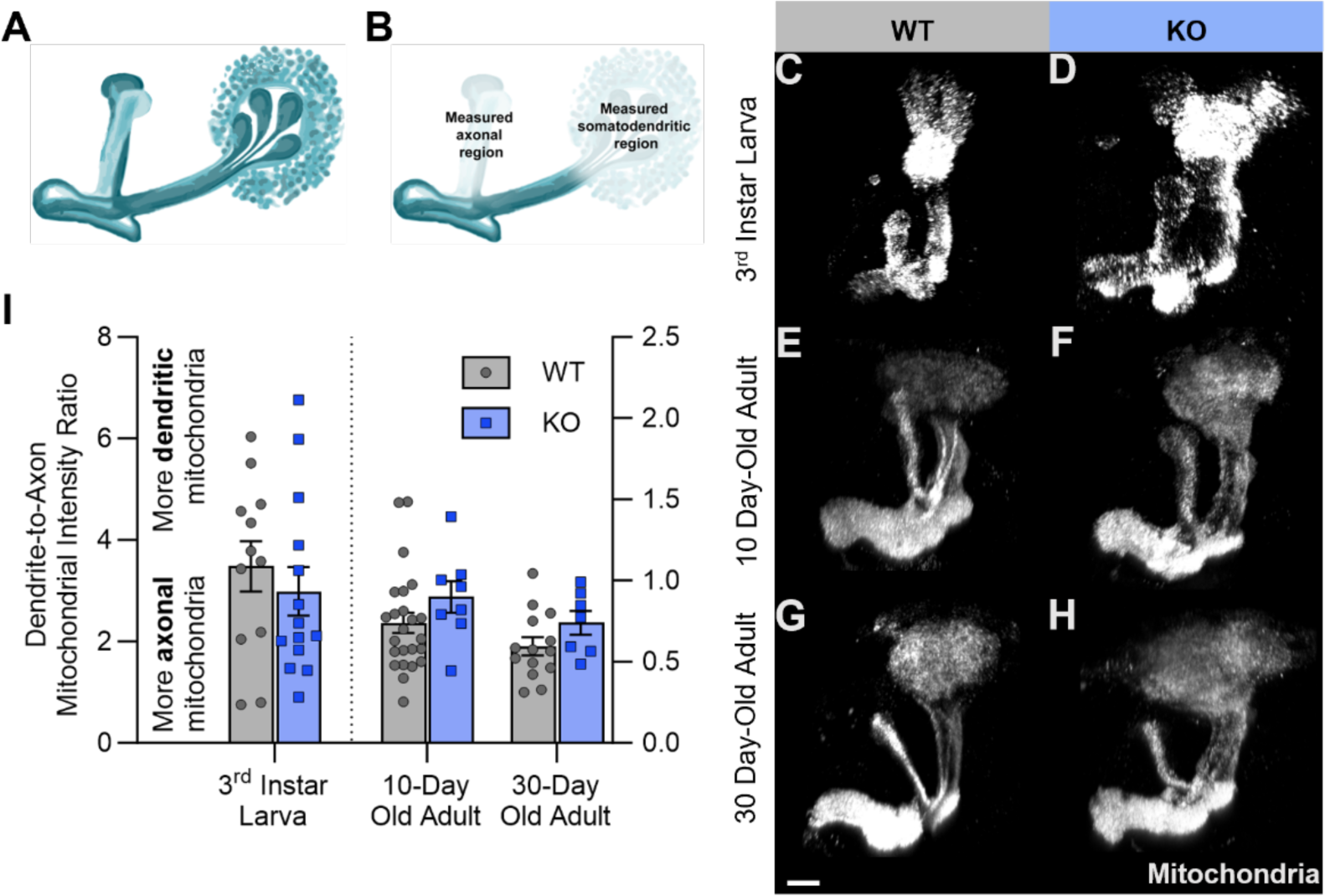
Cdk5α-KO does not cause gross disruption of mitochondrial distribution between somatodendritic and axonal compartments of larval and adult MB neurons. A) Schematic showing an oblique view of the MB in one hemisphere of the *Drosophila* brain. The calyx region containing the soma and dendrites is on the right and the bifurcated axons are on the left. Adapted from Armstrong et al., 1998 (with permission). B) MB schematic, as above, highlighting the regions used to quantify mitochondrial fluorescence in the axonal and somatodendritic compartments. To facilitate consistent measurement, the dorsal/alpha lobe (left) was measured for the axonal contribution as it was easier to consistently isolate in all the images. The MB calyx (right) was measured for the somatodendritic contribution. C-H) Representative normalized projections of mitochondrial fluorescence intensity in the MBs of WT 3^rd^ instar larvae (C), Cdk5α-KO 3^rd^ instar larvae (D), WT 10 day-old adults (E), Cdk5α-KO 10 day-old adults (F), WT 30 day-old adults (G), and Cdk5α-KO 30 day-old adults (H). Scale bar in lower left corner of the figure pane represents 20 µm. I) Quantification of the ratio of mitochondrial fluorescence intensity in the somatodendritic region to that of the axonal region of MB neurons of WT and Cdk5α-KO 3^rd^ instar larvae, 10 day-old adults, and 30 day-old adults. Larval groups compared using unpaired student’s *t*-test and adult groups compared using two-way ANOVA with Šidák’s multiple comparison Test. *p* values: * < .05; ** < .01, *** < .001, **** < .0001. Raw data associated with this figure and all subsequent figures can be found in Supplemental Table S1.

### 2.2. Mitochondria are selectively excluded from the AIS region of MB neurons in WT flies, and Cdk5α-KO disrupts this distribution

When examining mitochondrial distribution between the somatodendritic and axonal neuronal compartments of the MB, we observed a region of the proximal axon that appeared to lack mitochondrial fluorescence signal in WT flies. This region of the proximal axon seemed to overlap the area previously determined to correspond to the AIS of these neurons (Spurrier et al., 2019; Trunova et al., 2011). Interestingly, however, mitochondria were not excluded from this domain in Cdk5α-KO. To verify and better characterize this observation, we examined the mitochondrial fluorescence intensity through the region of the MB AIS and compared it to the pattern of other markers of the AIS boundaries. In MB neurons, the ubiquitous cytoskeletal protein actin can be used as a negative marker of the AIS owing to its relative exclusion from this domain (Spurrier et al., 2019; Trunova et al., 2011). Consistent with this, in WT 3^rd^ instar larvae, the actin-depleted domain correlates closely with the region lacking mitochondrial signal (Fig. 2A). In Cdk5α-KO 3^rd^ instar larvae, in contrast, there is no domain of mitochondrial exclusion at the normal position of the AIS (Fig. 2B). We also observe a similar mitochondrial distribution phenotype at the AIS in adult MB neurons as defined both by actin and by the cell surface protein Fasciclin II (FasII), an axon-specific protein that can be used to help identify the border between the AIS and the axonal compartment in MB γ-neurons (since the distal border of the AIS in adults can be difficult to visualize based solely on actin distribution; Spurrier et al., 2019; Trunova et al., 2011). Specifically, in adult WT flies, mitochondria are partially excluded from the AIS, with the proximal portion of the AIS showing striking depletion of mitochondria but with a progressive increase of mitochondrial signal towards the distal AIS (Figs. C-D).

**Figure 2.**
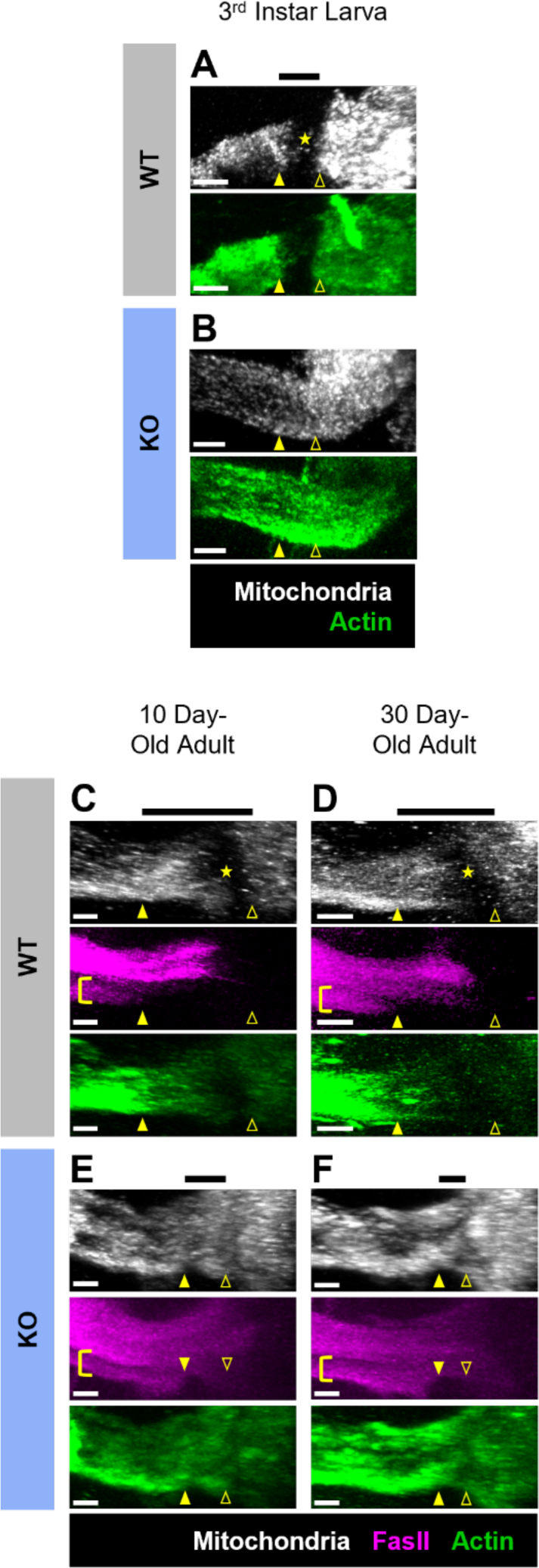
Mitochondria are depleted or excluded from the AIS region of MB neurons in WT flies, and Cdk5α-KO disrupts this mitochondrial distribution pattern. A-B) Representative normalized projections of mitochondrial and actin fluorescence intensity in MBs of WT 3^rd^ instar larvae (A) and Cdk5α-KO 3^rd^ instar larvae (B). The images are oriented such that the distal axons are to the left and the calyxes are to the right in these images and all MB images shown below. Yellow closed arrowheads indicate the distal border of the AIS and yellow open arrowheads indicate the proximal border as determined by the pattern of actin distribution in the WT image, or the approximate equivalent position in the Cdk5α-KO image based on morphological criteria (the emergence of bundled axons from the MB calyx). Yellow star in (A) indicates the region of mitochondrial exclusion in WT; note that in Cdk5α-KO there is not a region of mitochondrial exclusion. The black bar above (A) represents the length and position of the AIS as determined by the actin distribution; note that in Cdk5α-KO the AIS is completely ablated and thus there is no corresponding black bar for these images. White scale bars in the lower left corner of images represents 10 µm. C-F) Representative normalized projections of mitochondrial, FasII, and actin fluorescence intensity in MBs of WT 10 day-old adults (C), WT 30 day-old adults (D), Cdk5α-KO 10 day-old adults (E), and Cdk5α-KO 30 day-old adults (F). Yellow closed arrowheads indicate the distal border of the AIS as determined by the pattern of actin and FasII distribution as well as morphological criteria. Yellow brackets highlight the MB γ-neurons in which FasII demarcates the distal border of the AIS; note that FasII is not excluded from the AIS in other classes of MB neurons (α/β neurons). Yellow open arrowheads indicate the proximal border of the AIS as determined by the pattern of actin distribution in WT and by morphological criteria in both WT and Cdk5α-KO. Yellow stars indicate the region of maximal mitochondrial exclusion in the proximal portion of the AIS in WT (C-D). Note that in Cdk5α-KO images (E-F) the AIS is greatly reduced in size but not entirely absent; nonetheless, no region of complete mitochondrial exclusion is detected. Black bars above images represent the length and position of the AIS. White scale bars in the lower left corner of images represents 10 µm.

Similar to the observations in Cdk5α-KO 3^rd^ instar larvae, Cdk5α-KO adults also show a loss of mitochondrial exclusion within the AIS domain (Fig. 2E-F). Quantifying mitochondrial fluorophore intensity across the AIS region confirmed that there is a region of mitochondrial exclusion within the AIS in WT flies and that this exclusion is lost in Cdk5α-KO both in larvae and in adult flies (Fig. 3A-D). Notably, there is a striking depletion of mitochondria throughout the AIS in WT larvae and preferential depletion of mitochondrial from the proximal AIS in WT adults, but neither exclusion zone is observed upon fluorescence quantification in Cdk5a-KO (Fig. 3B-C). We do not observe any age-dependent change in the mitochondrial pattern at the AIS in either adult WT or Cdk5α-KO flies (Fig. 3D).

**Figure 3.**
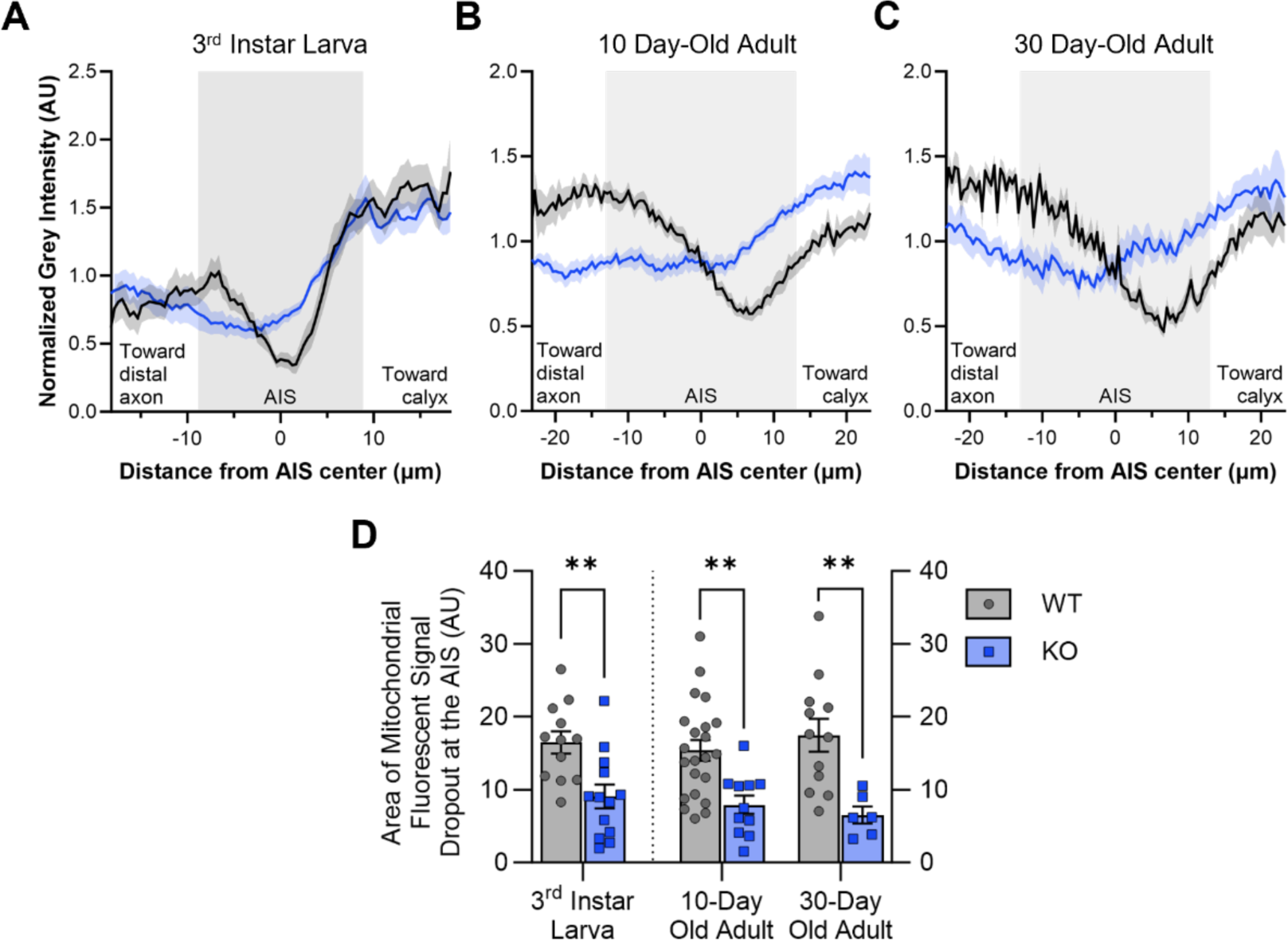
Quantification of mitochondrial fluorescence shows that mitochondria are partially or wholly excluded from the AIS region of WT MB neurons, and Cdk5α-KO disrupts this mitochondrial distribution pattern. A-C) Quantification of mitochondrial fluorescence intensity across the AIS region of MB neurons of WT and Cdk5α-KO 3^rd^ instar larvae (A), 10 day-old adults (B), and 30 day-old adults (C). Colored lines (black = WT; blue = KO) represent average normalized mitochondrial intensity value while colored shaded regions represents the standard error of the mean (SEM). The shaded background of the graph represents the average AIS size for larvae or adult, respectively, as determined by Trunova et al., 2011. More negative x-axis values are oriented towards the MB distal axon while more positive x-axis values are oriented towards the MB calyx. Sample sizes for 3^rd^ instar larvae: WT = 12, KO = 13. Sample sizes for 10 day-old adults: WT = 24, KO = 11. Sample sizes for 30 day-old adults: WT = 14, KO = 6. D) Quantification of the normalized, integrated mitochondrial intensity within the AIS region from Fig. 3A-C. Graphs show mean +/- SEM. Larval groups compared by unpaired student’s *t*-test and adult groups compared by two-way ANOVA with Šidák’s multiple comparison test. *p* values: * < .05; ** < .01, *** < .001, **** < .0001.

### 2.3. Structural disruption of the AIS independent of Cdk5α-KO alters mitochondrial distribution in the somatodendritic, axonal, and AIS compartments of MB neurons

The loss of mitochondrial exclusion from the AIS region in MB neurons of Cdk5α-KO flies raises the question whether the observed changes in mitochondrial distribution in Cdk5α-KO are due directly to alterations of Cdk5α or are secondary to perturbation of AIS structure. To determine if perturbing the AIS is sufficient to cause loss of the AIS-associated zone of mitochondrial exclusion, we examined mitochondrial distribution in flies that lack a functional AIS due to the expression of a dominant-interfering fragment of the Ank2 gene, called Ank2-L4 (Jegla et al., 2016; Pielage et al., 2008). In a previous study, expression of this dominant-negative variant shortened the AIS or ablated it altogether from early developmental stages (at least by the 3^rd^ larval instar) as assayed by several well-characterized molecular markers (Spurrier et al., 2019). We find here, however, that structural disruption of the AIS by expression of Ank2-L4 does not alter the pattern of mitochondrial distribution at the AIS by a statistically significant amount in 3^rd^ instar larvae nor in 10 day-old adult flies (Figs. 4, 5). In contrast, in 30 day-old adult flies expressing Ank2-L4, there is significant mitochondrial redistribution into the AIS of MB neurons (Figs. 4, 5). This corresponds to the stage at which chronic expression of Ank2-L4 induces blebbing, swelling, and fragmentation of the proximal axon and at which MB cell numbers begin to decline (Spurrier et al., 2019). In addition, by comparing the mitochondrial fluorophore intensity between the somatodendritic and axonal compartments, we find that disruption of the AIS by expressing Ank2-L4 is associated with an increase in the ratio of somatodendritic to axonal mitochondria in adult flies but not in larvae (Fig. 6).

**Figure 4.**
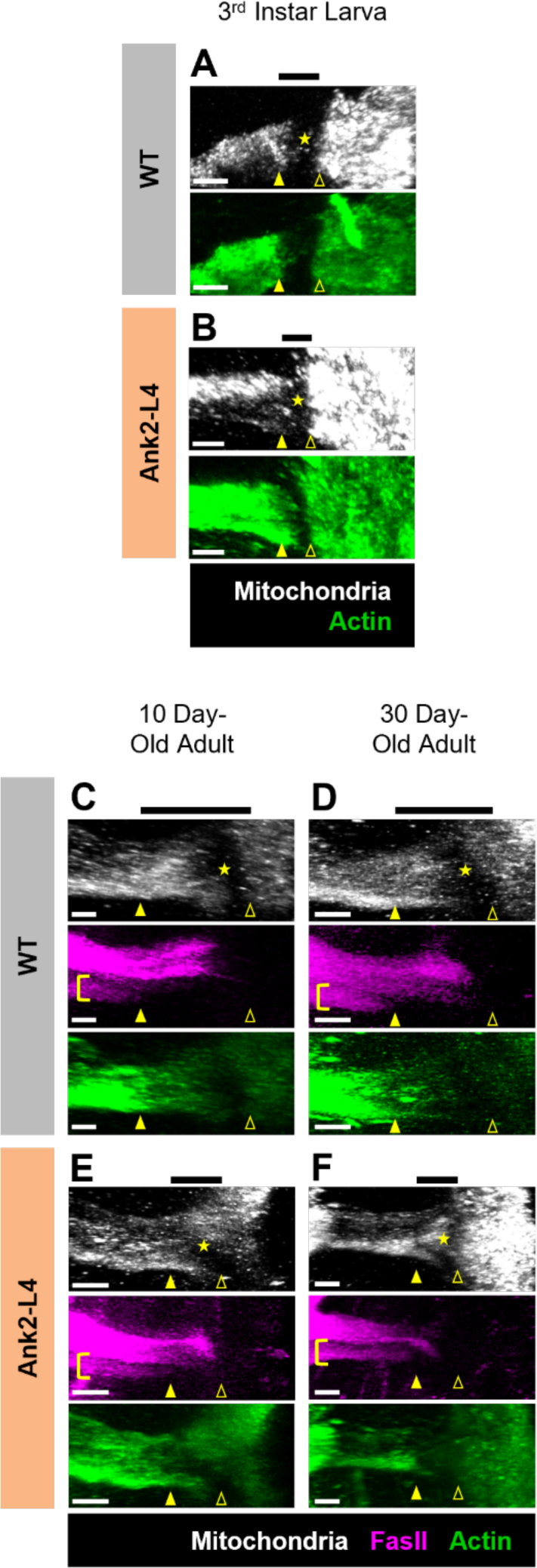
Structural disruption of the AIS by Ank2-L4 expression alters mitochondrial distribution in the AIS compartment of MB neurons only in old adults. A-B) Representative normalized projections of mitochondrial and actin fluorescence intensity in MBs of WT 3^rd^ instar larvae (A) and Ank2-L4 3^rd^ instar larvae (B). Yellow closed arrowheads indicate the distal border of the AIS and yellow open arrowheads indicate the proximal border of the AIS as determined by the pattern of actin distribution. Yellow stars indicate the region of mitochondrial exclusion. Black bars above images represent the length and position of the AIS as determined by the pattern actin distribution. White scale bars in the lower left corner of images represents 10 µm. Note that WT images are the same as those used in Fig. 2A. C-F) Representative normalized projections of mitochondrial, FasII, and actin fluorescence intensity in MBs of WT 10 day-old adults (C), WT 30 day-old adults (D), Cdk5α-KO 10 day-old adults (E), and Cdk5α-KO 30 day-old adults (F). Yellow closed arrowheads indicate the distal border of the AIS and yellow open arrowheads indicate the proximal border of the AIS as determined by the pattern of actin and FasII distribution as well as morphological criteria. Yellow brackets highlight the MB γ-neurons in which FasII demarcates the distal border of the AIS. Yellow stars indicate the region of maximal mitochondrial exclusion. Black bars above images represent the length of the AIS. White scale bars in the lower left corner of images represents 10 µm. Note that WT images are the same as those used in Fig. 2C-D.

**Figure 5.**
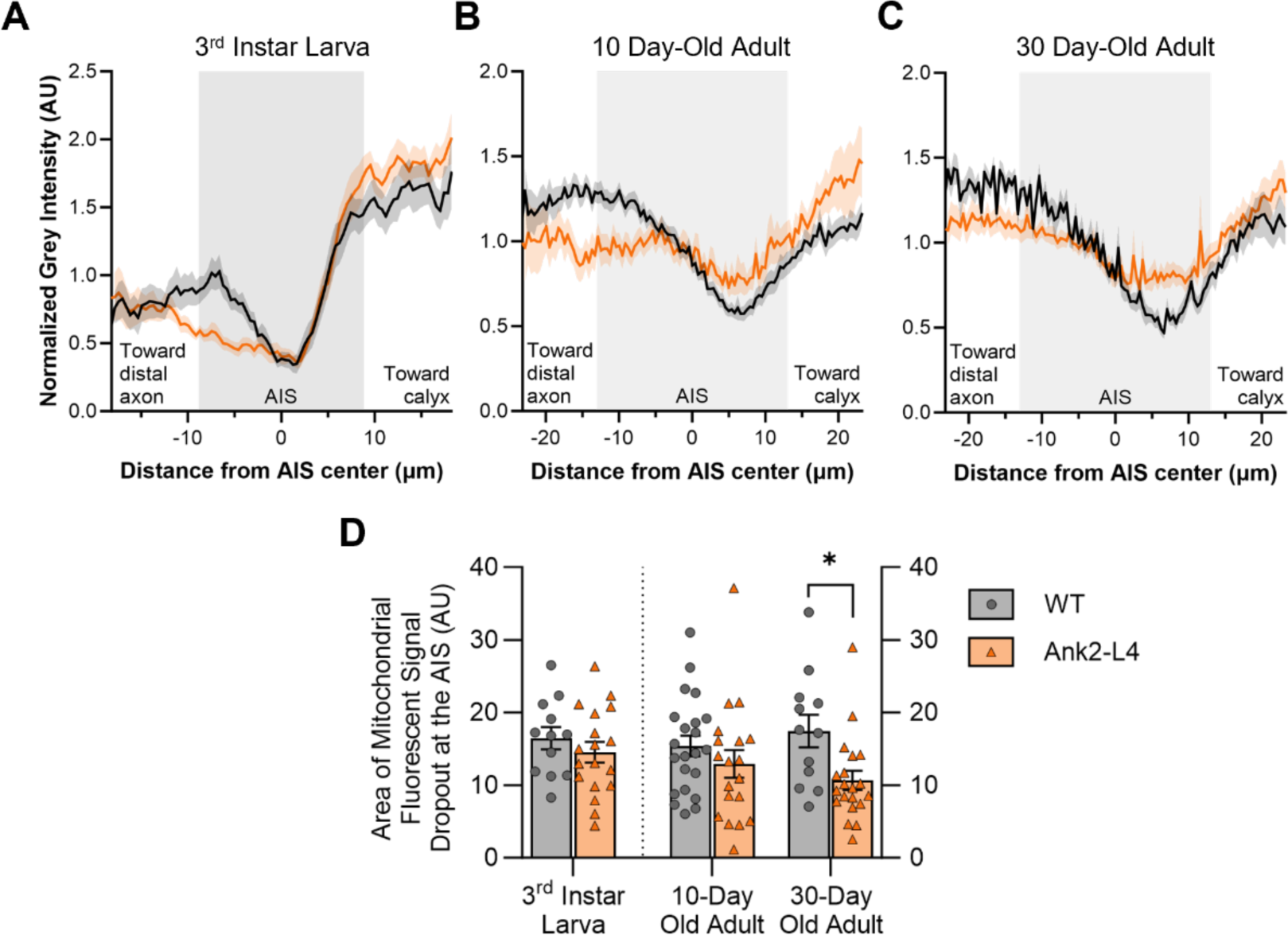
Quantification of mitochondrial fluorescence at the AIS shows that expression of Ank2-L4 causes mitochondrial redistribution into the AIS only in 30 day-old adults. A-C) Quantification of mitochondrial fluorescence intensity across the AIS region of MB neurons of WT and Ank2-L4 3^rd^ instar larvae (A), 10 day-old adults (B), and 30 day-old adults (C). Colored lines (black = WT; orange = Ank2-L4) represent average normalized mitochondrial intensity value while colored shaded regions represents the standard error of the mean (SEM). The shaded background of the graph represents the average AIS size for larvae or adult, respectively, as determined by Trunova et al., 2011. More negative x-axis values are oriented towards the MB distal axon while more positive x-axis values are oriented towards the MB calyx. Sample sizes for 3^rd^ instar larvae: WT = 12, Ank2-L4 = 18. Sample sizes for 10 day- old adults: WT = 24, Ank2-L4 = 19. Sample sizes for 30 day-old adults: WT = 14, Ank2-L4 = 20. D) Quantification of the normalized, integrated mitochondrial intensity within the AIS region from Fig. 5A-C. Graphs show mean +/- SEM. Larval groups compared by unpaired student’s *t*-test and adult groups compared by two-way ANOVA with Šidák’s multiple comparison test. *p* values: * < .05; ** < .01, *** < .001, **** < .0001.

**Figure 6.**
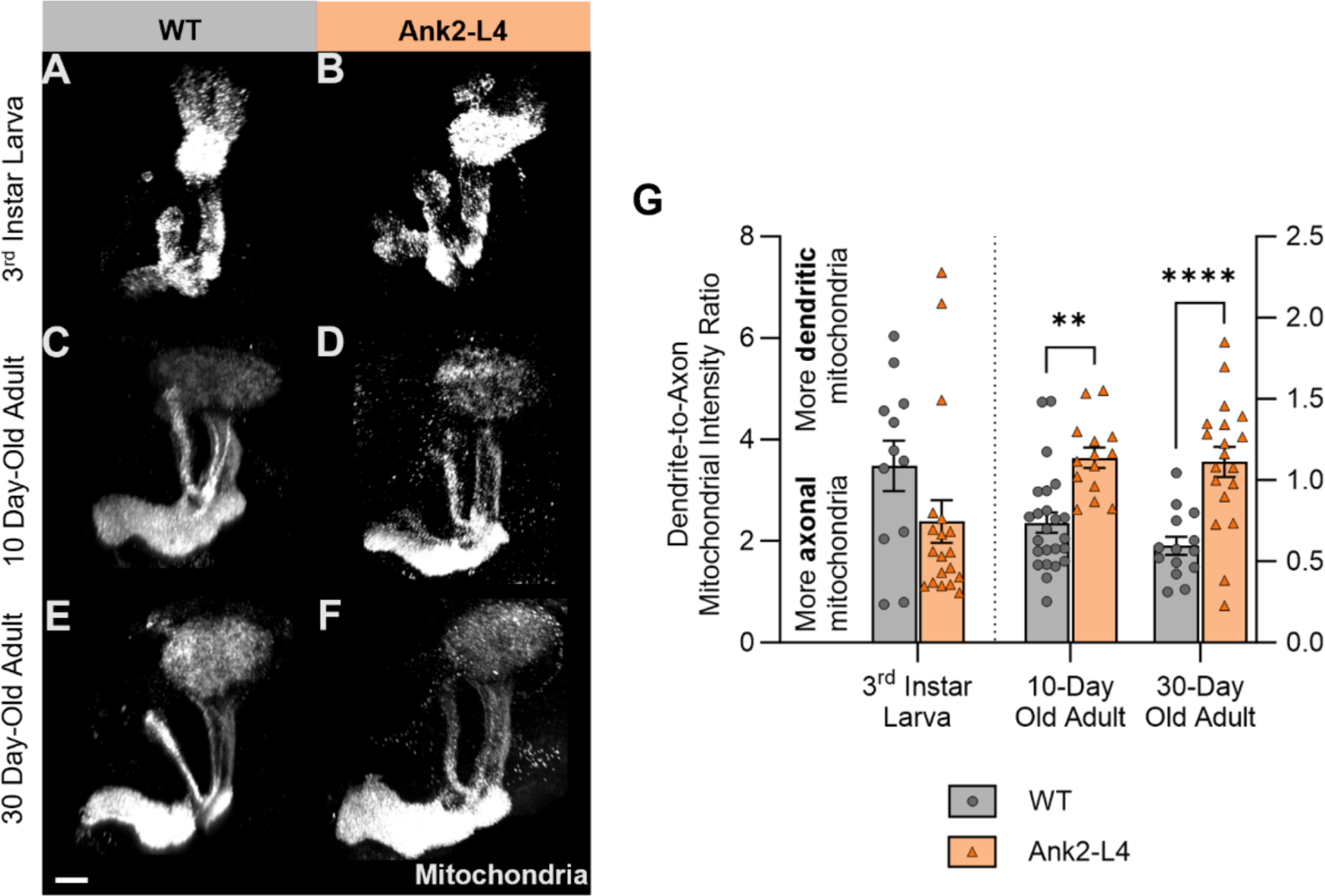
Ank2-L4 expression increases the ratio of somatodendritic to axonal mitochondria in 10 day-old and 30 day-old adult MB neurons. A-F) Representative normalized projections of mitochondrial fluorescence intensity in the MBs of WT 3^rd^ instar larvae (A), Ank2-L4 3^rd^ instar larvae (B), WT 10 day-old adults (C), Ank2-L4 10 day-old adults (D), WT 30 day-old adults (E), and Ank2-L4 30 day-old adults (F). Scale bar in lower left corner of the figure pane represents 20 µm. Note that WT images are the same as those used in Fig. 1. G) Quantification of the ratio of mitochondrial fluorescence intensity in the somatodendritic region to that of the axonal region of MB neurons of WT and Ank2-L4 3^rd^ instar larvae, 10 day-old adults, and 30 day-old adults. Larval groups compared using Mann-Whitney *U* test (Kolmogorov-Smirnov normality test, *p* < .05) and adult groups compared using two-way ANOVA with Šidák’s multiple comparison test. *p* values: * < .05; ** < .01, *** < .001, **** < .0001.

## 3. DISCUSSION

Here, we find that disruption of the AIS can be associated with gross alterations of mitochondrial distribution as neurons expressing a dominant-negative fragment of Ank2, Ank2-L4, display a relative overabundance of somatodendritic mitochondria relative to axonal mitochondria in adulthood, though not during development. On the contrary, perturbing the AIS by Cdk5α-KO does not shift the distribution of mitochondria between the somatodendritic and axonal compartments in larval or adult flies, excluding such gross redistribution as a potential mechanism of Cdk5-associated neurodegeneration. We also find that mitochondrial exclusion from the AIS is a feature of healthy neurons in the *Drosophila* MB, extending to *in vivo* central nervous system neurons observations made previously in peripheral neurons *in vivo* and cortical neurons *in vitro* (Tamada et al., 2021; Tjiang & Zempel, 2022). Examining mitochondrial distribution at the AIS further reveals that in Cdk5α-KO, a condition known to cause age-dependent axonal degeneration and subsequent neuron cell loss, mitochondria redistribute into the AIS region starting in development and persisting through adulthood, concomitant with loss of the AIS domain. Expression of Ank2-L4 also causes severe shortening or loss of the AIS throughout development and adulthood and leads to age-dependent axonal degeneration and neurodegeneration, but it does not cause mitochondria to redistribute into the AIS region until late adulthood, concomitant with the onset of age-dependent pathology. These findings suggest, first, that mitochondrial redistribution into the AIS domain in Cdk5α- KO is not a trivial consequence of disrupted AIS structure, but rather that it is likely to be a more direct consequence of altered Cdk5 activity since AIS structure can be disrupted directly by Ank2-L4 without triggering immediate mitochondrial redistribution. Second, they suggest that mitochondrial redistribution into the AIS is not by itself sufficient to initiate immediate degeneration since MB degeneration is delayed by more than 30 days after mitochondrial redistribution into the AIS in Cdk5α-KO. These data do not, however, exclude the possibility that mitochondrial accumulation in the AIS may be associated with late steps in axonal degeneration since such accumulation is observed in two mechanistically distinct neurodegenerative processes. Finally, while we find that the ratio of somatodendritic to axonal mitochondria becomes altered in concert with AIS disruption upon expression of Ank2-L4, we cannot distinguish whether these two effects are parallel consequences of Ank2-L4 or whether AIS disruption in this context is itself responsible for the change in overall mitochondrial distribution.

It is interesting that mitochondrial distribution may offer a unique tool with which to query the organization and properties of the AIS. While previous studies have shown strong concordance in the distribution of multiple AIS markers upon application of various manipulations (Cruz et al., 2009; Jegla et al., 2016; Marin et al., 2016; Spurrier et al., 2019; Trunova et al., 2011), here we find that the distribution of mitochondria in and around the AIS follows a somewhat different pattern. Specifically, mitochondria redistribute to the shrunken borders of the AIS from early development in Cdk5α-KO, but, upon expression of Ank2-L4, mitochondria only redistribute to the shrunken borders of the AIS at late adult stages, concomitant with the onset of overt degeneration. Previous experiments had provided evidence that the genetic pathways leading to AIS alteration by Cdk5α-KO and Ank2-L4 are distinct, but those experiments failed to identify a molecular pathway that accounted for the difference (Spurrier et al., 2019). The data here suggest that mitochondrial distribution may be an effective assay to discriminate these pathways in the future.

Numerous studies have reported accumulation of mitochondria, as well as other molecular cargo, within the AIS in various pathological contexts (Chang et al., 2006; Hirano et al., 1984; Sasaki et al., 2005; Sasaki & Iwata, 1996, 2007). One interpretation of the evidence posits that this accumulation of cargo represents a pathologic “traffic jam” preventing passage of molecular cargo between the axonal and somatodendritic compartments (Sabharwal & Koushika, 2019). However, our data suggest that accumulation of mitochondria in the AIS does not necessarily impair mitochondrial distribution to the distal axon. In fact, this accumulation of mitochondria may be part of a physiological damage repair mechanism that occurs in conjunction with the temporary dismantling of the AIS after injury (Kiryu-Seo et al., 2022; Tamada et al., 2021; Tjiang & Zempel, 2022). Our data suggest that similar processes may occur in the context of chronic neurodegenerative insults and raise a question about whether the redistribution of mitochondria into the AIS region may actually be cytoprotective. Although reasons for mitochondrial redistribution into the AIS in the context of injury and disease are becoming more evident, it remains unclear why mitochondria are seemingly excluded from the energetically demanding AIS under normal circumstances (Yang et al., 2023).

Our data also suggest that mitochondrial distribution within the AIS region is regulated independently from mitochondrial transport through the AIS. The factors governing mitochondrial distribution may rely upon functional parameters such as mitochondria-calcium interactions (Saotome et al., 2008; Yi et al., 2004; Zhang et al., 2010) or upon structural factors such as actin patches (Balasanyan et al., 2017; Watanabe et al., 2012) or syntaphilin anchors (Chen et al., 2009; Chen & Sheng, 2013; Kang et al., 2008) akin to mechanisms that occur at the nodes of Ranvier. However, further study is required to determine precisely which factors anchor mitochondria outside the AIS and which factors permit mitochondria to pass through the AIS. Importantly, our data indicate that age and disease can alter the distribution of mitochondria at the AIS and, thus, suggest that future experiments should investigate how the factors controlling this distribution may be modulated by aging and disease.

One intriguing possibility is that one of these factors that modifies mitochondrial localization within the AIS region may be the neurodegeneration-associated protein tau. Tau is known to play multiple roles at the AIS, including in activity-dependent plasticity, structural integrity, and selective exclusion of molecular cargo (Hatch et al., 2017; Li et al., 2011; Sohn et al., 2019). In the context of disease, however, the ability of tau mutants or hyperphosphorylated tau to interact with their molecular targets, including microtubules and mitochondrial proteins, is impaired (Tracy et al., 2022). It is intriguing to speculate that altered tau function could explain the differences in mitochondrial redistribution at the AIS between Cdk5α-KO and Ank2-L4 expression. Altering the activity of Cdk5, which is known to lead to the hyperphosphorylation of tau, causes mitochondrial redistribution into the AIS throughout the lifespan while expression of Ank2-L4, which is not known to directly cause changes in tau phosphorylation, only shows redistribution of mitochondria into the AIS at older ages when tau is known to undergo age- dependent hyperphosphorylation (Higham et al., 2019; Pao & Tsai, 2021). It is conceivable, therefore, that impaired interactions between tau and mitochondria may underlie the redistribution of mitochondria into the AIS, but more study will be required to determine if such a mechanism exists.

## 4. METHODS

### 4.1. Fly stocks, maintenance, and aging

All flies were maintained on a standard cornmeal-molasses *Drosophila* media (Caltech media; KD Medical, Columbia, MD) at 25°C and 50% humidity on a 12:12 hour light:dark cycle. All experiments were performed in male wandering 3^rd^ instar larvae and male adult flies in an Oregon Red (w^+^) background. The stocks used in these experiments were as follows: *201Y-GAL4* (BDSC, #4440), *UAS-DsRed.mito 47A* (BDSC, #93056), *UAS-myc-Actin* (a gift from Cheng-Ting Chien, Academia Sinica, Taipei, Taiwan), and *UAS-Venus- Ank2-L4* (a gift from Jan Pielage, RPTU, Landau, Germany). The Cdk5α-KO condition (*w^+^; Cdk5α ^20C^/Df(Cdk5α)^C2^; +*) has been described in detail elsewhere (Connell-Crowley et al., 2000, 2007; Spurrier et al., 2018; Trunova & Giniger, 2012).

For the assessment of the mitochondrial distribution in MB neurons, the γ-neuron-specific *GAL4* driver *201Y-GAL4* was used to express *UAS-DsRed.mito 47A* and *UAS-myc-Actin* in WT, Cdk5α-KO, and Ank2-L4 conditions. The resulting experimental genotypes are as follows: WT (*w+; 201Y-GAL4/+; UAS-DsRed.mito 47A, UAS-myc-Actin/+*), Cdk5α-KO (*w+; Df(Cdk5α)^C2^, 201Y-GAL4/Cdk5α^20C^; UAS-DsRed.mito 47A, UAS-myc-Actin/+*), and Ank2-L4 (*w+; 201Y-GAL4/+; UAS-DsRed.mito 47A, UAS-myc-Actin/UAS-Venus- Ank2-L4*).

For aging of adult flies, males and females were collected within 24 hours of eclosion and transferred to fresh vials. After three days, males were collected and placed into fresh vials. Flies were flipped to fresh vials twice weekly while aging until they reached the appropriate experimental age.

### 4.2. Immunohistochemistry

For immunohistochemistry of larval brains, brains were dissected from male wandering 3^rd^ instar larvae in ice-cold 1x PBS. Brains were subsequently fixed in 4% paraformaldehyde for 25 minutes at room temperature. Brains were post-fixed in 4% paraformaldehyde plus 0.5% Triton X-100 for 25 minutes at room temperature. After three washes in PBS with 0.5% Triton X-100 (PBST), brains were blocked for 2 hours in blocking buffer (PBST supplemented with 4% bovine serum albumin, 4% normal goat serum, and 4% normal donkey serum) at room temperature. Brains were incubated in mouse anti-FasII (1:200, Developmental Studies Hybridoma Bank) and rabbit anti-myc (1:1000, Sigma-Aldrich) diluted in blocking buffer for 16 hours at 4°C. The rabbit anti-myc antibody was preadsorbed with Drosophila embryos prior to use. Brains were then washed thrice in PBST at room temperature and incubated in Alexa Fluor secondary antibodies (1:500, ThermoFisher Scientific) diluted in blocking buffer for 4 hours at room temperature. Prior to mounting, brains were washed thrice with PBS at room temperature. To prevent squishing of the brains, brains were mounted on slides between two number 1 glass coverslip pieces before covering with VectaShield mounting medium (Vector Laboratories) and a coverslip.

For immunohistochemistry of adult brains, the procedure was the same as for larval brains except for the following incubation timing differences: fixation for 35 minutes, post-fixation for 35 minutes, primary antibody incubation for 48 hours, and secondary antibody incubation for 16 hours.

### 4.3. Confocal microscopy and image analysis

Images were acquired using a LSM880 confocal microscope (Zeiss) equipped with a 40X (1.2 NA) water objective. Each MB hemisphere was imaged as a Z-stack with a 1.0 µm step size covering approximately 120 µm. All microscope settings were identical for all brains imaged. All image analysis was done using ImageJ (version 1.53t, NIH).

To compare the mitochondrial fluorescence intensity between the somatodendritic and axonal neuronal compartments, all images were first converted to 32-bit format and thresholded to the same value with the values below the threshold set to “not a number”. Images were then rotated such that they were all in an identical orientation with both the calyx and the dorsal/alpha lobes in plane. A custom ROI was then drawn around the calyx of the MB – the calyx harbors the dendritic arbor as well as cell bodies of the MB neurons – and the summed intensity of the non-zero pixels and the area of the non-zero pixels was calculated. The average mitochondrial intensity in the MB calyx was calculated as the summed intensity divided by the non-zero area. This process was repeated to calculate the average mitochondrial intensity in the MB dorsal/alpha lobe. The dorsal/alpha lobe was used instead of all axonal lobes as it was easier to consistently isolate in the images. The somatodendritic-to-axonal mitochondrial intensity ratio was then computed by dividing the average mitochondrial intensity in the MB calyx by the average mitochondrial intensity in the MB dorsal/alpha lobe.

To measure the mitochondrial fluorescence intensity in the AIS region, images were first converted to 32-bit format, and all images were thresholded to the same value with the values below the threshold set to “not a number”. Images were then rotated such that they were in an identical orientation with the AIS region in plane. Image z-stacks were then flattened via the maximum fluorescence projection function in ImageJ. A rectangular ROI approximately 37 µm (for larvae) or 47 µm (for adults) in length and approximately the width of the MB peduncle was drawn across the AIS region with its center aligned manually with the approximate center of the AIS. The center of the AIS was approximated using the distribution patterns of the faithful AIS markers FasII and actin as well as morphological features of the tissue (the emergence of bundled axons from the MB calyx). The intensity histogram of this ROI was taken with a bin size of approximately 0.41 µm. Raw intensity values across the ROI were then normalized to the maximum intensity value for each image. The mean and standard error was then calculated for each genetic condition and age from each normalized bin intensity value. This normalized mean and standard error was then plotted across the length of the ROI (37 µm for larvae or 47 µm for adults).

To quantify the drop in normalized mitochondrial fluorescence intensity across the AIS (“signal dropout”), the normalized mitochondrial fluorescence intensity was plotted across the length of the ROI (37 µm for larvae or 47 µm for adults) for each image. A line was then drawn which intersected the normalized mitochondrial fluorescence intensity plot at the distal and proximal edges of the AIS (as determined by the faithful AIS markers actin and FasII as well as morphological features of the tissue). The area enclosed by this straight line and the normalized mitochondrial fluorescence intensity plot was then calculated using the ImageJ “Wand” tool and labeled as the area of mitochondrial signal dropout.

All image visualization was done using Imaris (version 9.5, Oxford Instruments) software. Z-stack images were projected using the 3D View tool. Images were then rotated to the desired orientation and cropped to remove out-of-plane signal. Due to differences in fluorescence intensity across genotypes, signal intensity was adjusted to allow for cross-sample visual comparisons.

### 4.4. Statistical analyses

All statistical analyses were performed using Prism (GraphPad, version 10). Data were tested for normality using the Kolmogorov-Smirnov test and for equality of variances using the F-test of equality of variances or Barlett’s test. Specific statistical tests used are described in the Figure Legends. All image analysis was completed blind to genotype and age. In all experiments, one biological replicate is represented by a single fly. When appropriate, measurements from each MB hemisphere from a single brain were treated as technical replicates, and these measurements were averaged together.

## ACKNOWLEDGEMENTS

We wish to thank all current and former members of our lab for helpful input that helped to shape the experiments discussed here. Specifically, we would like to thank Arvind Shukla, Andrew Scott, Rameen Forghani, Phil McQueen, Hitesh Chaouhan, and Suparna Saha. We are also thankful to Bill Rebeck, Tom Coate, Derek Narendra, and Tingting Wang for helpful insights, discussions, and suggestions during the course of these experiments and for comments on the manuscript. We are also grateful for the microscopy assistance of Stephen Wincovitch of the NHGRI Cytogenetics and Microscopy Core Facility. We are also thankful to Derek Narendra of the National Institute of Neurological Disorders and Stroke (NINDS) for sharing lab space and equipment. Reagents essential for these experiments were provided by the Developmental Studies Hybridoma Bank (University of Iowa), from the Bloomington Drosophila Stock Center (University of Indiana), and as generous gifts from C-T Chien and Jan Pielage. This work was supported by the Basic Neuroscience Program of the Intramural Research Program of the NINDS / National Institutes of Health (NIH) (Z01 NS003106 to E.G.). All authors report no conflicts of interest. All relevant data can be found within the article or in the supplementary information.

## AUTHOR CONTRIBUTIONS

APKW: conceptualization, methodology, validation, formal analysis, investigation, writing – original draft, writing – review & editing, visualization

BTH: writing – review & editing, supervision

EG: conceptualization, resources, writing – review & editing, supervision, funding acquisition

## Abbreviations

AIS: Axon initial segment
AnkG: Ankyrin G
Ank2: Ankyrin 2
Ank2-L4: Portion of the L4 exons of Ankyrin 2 corresponding to a fragment of the C-terminus of the Ank2 isoform Ank2-L
Cdk5α-KO: Knockout of Cdk5α
FasII: Fasciclin II
KO: Knockout of Cdk5α
MB: Mushroom body
WT: Wild-type

---

*Supplementary Table 1 Legend*

Raw data associated with Figs. 1, 3, 5-6

4.1. Supplementary Table 1

“Table S1_Raw Data.xlsx”

## Notes

### Competing Interest Statement

The authors have declared no competing interest.

